# A face identity hallucination (palinopsia) generated by intracerebral stimulation of the face-selective right lateral fusiform cortex

**DOI:** 10.1101/142786

**Authors:** Jacques Jonas, Hélène Brissart, Gabriela Hossu, Sophie Colnat-Coulbois, Jean-Pierre Vignal, Bruno Rossion, Louis Maillard

**Affiliations:** Service de Neurologie, Centre Hospitalier Universitaire de Nancy, 29 Avenue du Maréchal de Lattre de Tassigny, 54000 Nancy, France; CRAN, UMR 7039, CNRS et Université de Lorraine, Campus Sciences, Boulevard des Aiguillettes, 54500 Vandœuvre-lès-Nancy, France; Institut de recherche en sciences psychologiques, Université Catholique de Louvain, 10 Place du Cardinal Mercier, 1348 Louvain-La-Neuve, Belgium; CIC-IT, Centre Hospitalier Universitaire de Nancy, 5 Rue du Morvan, 54500 Vandœuvre-lès-Nancy, France; Service de Neurochirurgie, Centre Hospitalier Universitaire de Nancy, 29 Avenue du Maréchal de Lattre de Tassigny, 54000 Nancy, France

**Keywords:** intracerebral recordings, electrical brain stimulation, palinopsia, fusiform gyrus, fusiform face area

## Abstract

We report the case of a patient (MB, young female human subject) who systematically experiences confusion between perceived facial identities when electrically stimulated inside the lateral section of the right fusiform gyrus. In the presence of a face stimulus (an experimenter or a photograph), intracerebral electrical stimulation in this region generates a perceptual hallucination of an individual facial part integrated within the whole perceived face, i.e. facial palinopsia. In the presence of a distracting stimulus (visual scene or object picture), the patient also experiences an individual face percept superimposed on the non-face stimulus. The stimulation site evoking this category-selective transient palinopsia is localized in a region showing highly selective responses to faces both with functional magnetic resonance imaging (“Fusiform Face Area”, “FFA”) and intracerebral electrophysiological recordings during fast periodic visual stimulation (FPVS). Importantly, the largest electrophysiological response to fast periodic changes of facial identity is also found at this location. Altogether, these observations suggest that a local face-selective region of the right lateral fusiform gyrus suffices to generate a vivid percept of an individual face, supporting the active role of this region in individual face representation.

## 1. Introduction

Individual face recognition plays a critical role in human social interactions. Studies of patients showing individual face recognition impairment after brain damage (i.e., prosopagnosia, following Bodamer, 1947; see Della Sala and Young, 2003 for a description of the earliest case of Quaglino and Borelli in 1867) have long suggested that this brain function is supported by a large territory of the human ventral occipito-temporal cortex (VOTC), from the occipital pole to the temporal pole, with a right hemispheric advantage (Hécaen and Angelergues, 1962; Meadows, 1974; Sergent and Signoret, 1992; Barton, 2008; Rossion, 2014).

Within this cortical territory, the lateral section of the right posterior/middle fusiform gyrus (latFG) may be particularly important, as this region shows the largest selective response to faces both in neuroimaging (“Fusiform Face Area”, “FFA”, e.g., fMRI: Puce et al., 1995; Kanwisher et al., 1997; Kanwisher, 2017; PET: Sergent et al., 1992; Rossion et al., 2000) and intracerebral recordings (Jonas et al., 2016). Recent studies have shown that electrical intracranial stimulation over the right – but not the left – latFG may elicit transient impairment in face perception, with patients reporting a selective distortion of the visual face input (i.e., distortion of people’s faces in the room: “prosopometamorphopsia”, Parvizi et al., 2012; Rangarajan et al., 2014). Importantly, patients with prosopometamorphopsia, often a transient phenomenon observed shortly after brain damage (e.g., Bodamer, 1947: case 3/patient B), do not present with major difficulties in individual face recognition, in spite of their perceptual distortions (Bodamer, 1947; Hécaen and Angelergues, 1962; Nass et al., 1985; Trojano et al., 2009; Hwang et al, 2012). Although stimulation of the right latFG inducing prosopometamorphopsia offers a causal link between face perception and this area, impairment in individual face recognition, which characterizes patients with prosopagnosia (Hécaen and Angelergues, 1962; Sergent and Signoret, 1992; Rossion, 2014), has so far not been observed following electrical stimulation of this region.

Here, we report a rare case of confusion of facial identity following focal electrical stimulation directly in the grey matter of the right latFG, without any face distortion. When stimulated only in this region, the patient (MB) systematically reported visual hallucinations characterized by the recurrence of individual facial parts (facial palinopsia, Critchley, 1951) integrated within the whole perceived face (a person in the room, or a photograph). Importantly, the electrode contact evoking this category-selective transient palinopsia is localized in a region showing highly selective responses to faces both with functional magnetic resonance imaging and with intracerebral recordings. These observations suggest that a local face-selective region of the right latFG may suffice to generate a vivid hallucination of an individual face, highlighting the active role of this region in individual face representation.

## 2. Materials and Methods

### 2.1 Case description and neuropsychological assessment

The subject is a 30-year-old woman (MB) with refractory focal epilepsy. Intracerebral stereo-electroencephalography (SEEG) delineated her epileptogenic zone in the right parietal cortex. The patient was right-handed as attested by the Edinburgh Handedness Inventory (Oldfield, 1971). She gave written consent to participate in the experimental procedures, which were part of the clinical investigation.

MB showed a general intelligence level in the normal range (full-scale IQ of 97). Neuropsychological evaluations revealed normal performance on memory (Taylor Complex Figure, Selective Reminding Test), language (DO80 naming test) and basic visual perception (Visual Object and Space Perception battery, VOSP) functions. The patient never complained of individual face recognition difficulties in everyday life, nor during or after epileptic seizures. Before intracerebral implantation, we conducted an extensive series of behavioral tests to assess MB’s face/object perception and memory. Ten control participants (age-, sex- and education level-matched controls) performed the same tests. To compare the results of MB to the control participants, we used the modified t-test of Crawford-Howell for single-case studies (Crawford and Howell, 1998) with a p value of &0.05 considered as statistically significant. These tests included: (1) face/no face categorization test (Mooney faces, experiment 16 in Busigny et al., 2010); (2) tests of face individuation including the Benton Face Recognition Test (BFRT, Benton et al., 1983), the Cambridge Face Memory Test (CFMT, Duchaine and Nakayama, 2006), an individual face- and car-matching at upright and inverted orientations (experiment 4 in Busigny and Rossion, 2010), as well as an individual matching task of faces presented in different viewpoints (experiment 22 in Busigny et al., 2010); (3) tests of visual memory including an old/new face task (encoding phase followed by an old/new forced choice decision with faces, experiment 3 in Busigny et al., 2010) and an old/new bird task (same task with bird pictures, using the same parameters as for faces); (4) a famous face recognition test (CELEB test, Busigny et al., 2014).

The results of these tests are shown in Table 1. MB performed in the normal range compared to matched normal controls for nearly all tests, either in accuracy or in response times, except that she was significantly slower at a visual memory test with faces and birds (old/new face and old/new bird tests, see Table 1), a slowing down that is often found in epileptic patients under medication. Her decrease of performance for inverted compared to upright faces was also in the normal range in accuracy and response times (t=0.68, p=0.51 and t=73, p=0.48 respectively, revised standardized difference test, Crawford and Garthwaite, 2005). Taken together, these results show that MB has normal face perception and memory abilities.

**Table 1.** Performance of patient MB and 10 control participants in neuropsychological tests of face/object perception and memory (Acc: accuracy; RT: reaction times in ms; BFRT: Benton Face Recognition Test; CFMT: Cambridge Face Memory Test; FRI: Face Recognition Index; NAI: Name Access Index). *Indicates lower performance compared to matched normal controls (p<0.05)

|  |  | Subject MB | Controls (n=10) | t-test (Crawford-Howell) |
| --- | --- | --- | --- | --- |
| Mooney faces | Acc | 97.5% | 92.1 ±3.2 | $t=1.61$, $p=0.07$ |
| | RT | 1708 | 1243 ±635 | $t=0.7$, $p=0.25$ |
| BFRT | Acc | 44/54 | 44.9 ±2.5 | $t=-0.34$, $p=0.37$ |
| | RT | 456 | 315 ±158 | $t=0.85$, $p=0.21$ |
| CFMT | Acc | 52/72 | 53.6 ±7.6 | $t=-0.2$, $p=0.42$ |
| Face matching<br>(different viewpoints) | Acc | 86% | 81.90 ±8.16 | $t=0.48$, $p=0.32$ |
| | RT | 4518 | 2761 ±975 | $t=1.72$, $p=0.6$ |
| Face matching<br>(upright and inverted) | Acc upright | 97.2% | 93 ±4.9 | $t=0.81$, $p=0.22$ |
| | inverted | 77.8% | 79.4 ±7.8 | $t=-0.2$, $p=0.42$ |
| | RT upright | 1791 | 1712 ±658 | $t=0.11$, $p=0.46$ |
| | inverted | 2334 | 1979 ±624 | $t=0.39$, $p=0.35$ |
| Car matching<br>(upright and inverted) | Acc upright | 100% | 96.7 ±9.6 | $t=0.33$, $p=0.37$ |
| | Inverted | 100% | 94.2 ±11.1 | $t=0.5$, $p=0.31$ |
| | RT upright | 1485 | 1712 ±658 | $t=-0.33$, $p=0.37$ |
| | inverted | 1601 | 1979 ±624 | $t=-0.58$, $p=0.29$ |
| Old/New face | Acc | 96.7% | 94 ±3.1 | $t=0.83$, $p=0.21$ |
| | RT | 3859* | 1959 ±820 | $t=2.21$, $p=0.03$ |
| Old/New bird | Acc | 83.3 | 84 ±7.3 | $t=-0.09$, $p=0.46$ |
| | RT | 4279* | 2227 ±900 | $t=2.17$, $p=0.03$ |
| CELEB test | FRI | 82 | 86.9 ±13 | $t=-0.36$, $p=0.36$ |
| | NAI | 95.1 | 90.7 ±12 | $t=0.35$, $p=0.37$ |

### 2.2 Stereotactic placement of intracerebral electrodes

Intra-cerebral electrodes (Dixi Medical, Besangon, France) were stereotactically implanted in the patient’s brain in order to delineate the seizure onset zone (SEEG, Talairach and Bancaud, 1973; Cardinale et al., 2013; Salado et al., in press). The sites of electrode implantation were determined based on non-invasive data collected during an earlier phase of the investigation. Each intracerebral electrode consists of a cylinder of 0.8 mm diameter and contains 8-15 contiguous contacts of 2 mm in length separated by 1.5 mm from edge to edge. A few days before surgery, a non-stereotactic T1 weighted MRI with gadolinium was carried out and imported into a computer-assisted software (Iplanstereotaxy, Brainlab, Germany). Each electrode trajectory was then determined according to the investigation planning with careful avoidance of vascular structures. The day of surgery, after induction of general anesthesia, the stereotactic frame (Leksell G-frame, Elekta, Sweden) was positioned on the patient’s head. A stereotactic CT-scan was then carried out and fused to the pre-operative non stereotactic MR. Stereotactic coordinates were then calculated for each trajectory. A post-operative non-stereotactic CT-scan was carried out and fused with a T1-weighted MRI to determine the exact position of each electrode.

The SEEG exploration took place in April 2014. Eleven electrodes were implanted in the right hemisphere targeting the occipito-parieto-temporal cortex and the hippocampus (Figure 1A). In total, these electrodes contained 138 individual recording contacts. Electrode F (containing 10 contacts, F1 to F10) was located in the right ventral temporal cortex, targeting specifically the latFG, the occipitotemporal sulcus and the inferior temporal gyrus (Figure 1B). No electrodes were placed in the left hemisphere.

**Figure 1.**
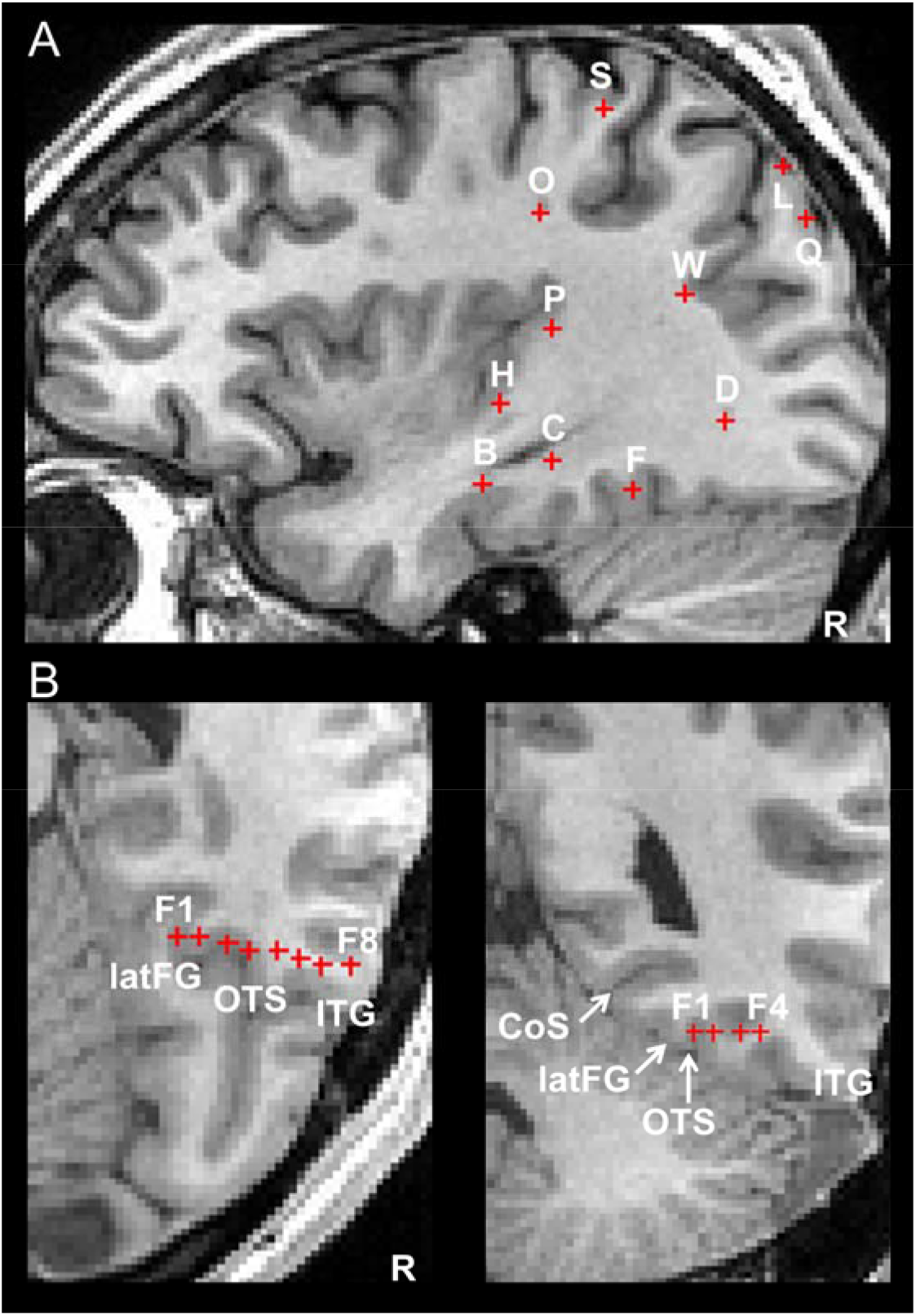
Locations of intracerebral electrodes implanted in subject MB’s brain. Each electrode is an array of individual recording contacts. Here, each contact is represented by a red cross. **A.** Locations of all electrodes on a sagittal slice. Since electrodes are usually implanted perpendicularly to the skull, only one contact per electrode is visible on a sagittal slice. **B.** Location of electrode F in the ventral temporal cortex on axial and coronal slices. Acronyms: CoS: collateral sulcus; ITG: inferior temporal sulcus; latFG: lateral fusiform gyrus; OTS: occipito-temporal sulcus.

The SEEG signal was recorded at a 512 kHz sampling rate on a 256-channel amplifier (4 SD LTM 64 Headbox, Micromed, Italy). The reference electrode during data acquisition was a midline prefrontal scalp electrode (FPz).

### 2.3 Intracerebral electrical stimulations

#### 2.3.1 General procedure

Electrical intracerebral stimulation of the right occipito-parieto-tempral cortex and hippocampus was carried out while the patient performed active recognition or passive viewing of visual objects (photographs of famous face, famous scenes, common objects, unknown faces or real faces, Table 2). These stimulations were applied between two contiguous contacts on the same electrode and performed at 50 Hz during 5s at low intensities ranging from 1 to 1.6 mA (i.e., usual stimulation settings in SEEG). MB was not aware of the stimulation onset and termination, the stimulation site and the nature of the effects that could be potentially elicited. In total, 59 electrical stimulations were performed (Table 2). Authors JJ, JPV, LM and BR were present during electrical stimulations, which were video recorded.

**Table 2.** Number of electrical stimulations performed and type of task required. Acronyms: CoS: collateral sulcus; IOG: inferior occipital gyrus; ITG: inferior temporal gyrus; latFG: lateral fusiform gyrus; LG: lingual gyrus; MTG: middle temporal gyrus; OTS: occipito-temporal sulcus; STS: superior temporal gyrus.

|  | Active recognition |  |  | Passive viewing |  |  |
| --- | --- | --- | --- | --- | --- | --- |
|  | Famous faces | Famous scenes | Common objects | Unknown faces | Common objects | Real faces |
| Contacts F1-F2 (latFG, OTS) | 3 | 1 | 1 | 1 | 1 | 2 |
| Ventral temporal cortex except contacts F1-F2 (OTS, CoS) | 3 |  | 1 |  |  | 2 |
| Lateral temporal cortex (ITG, MTG, STG) | 4 |  | 2 |  | 1 | 2 |
| Occipital cortex (IOG, lateral occipital cortex, cuneus, LG) | 3 | 1 |  |  | 4 | 3 |
| Parietal cortex (lateral parietal cortex, precuneus) | 4 | 2 | 3 |  | 2 | 11 |
| Hippocampus |  |  | 2 |  |  |  |
| Total | 17 | 4 | 9 | 1 | 8 | 20 |

#### 2.3.2 Active recognition

Thirty stimulations were carried out during the recognition of sets of photographs of the same category presented one by one (famous faces without external features, common objects or famous scenes). MB had to name each photograph in turn. For each set, the stimulation was triggered randomly during the presentation of 1 or 2 successive photographs. We only used photographs that were easily recognized by MB before the stimulation procedure.

#### 2.3.3 Passive viewing

Twenty-nine stimulations were carried out while MB was asked to passively watch photographs of unknown faces, common objects, or real faces of people in the room. MB was instructed to describe any perceptual changes she experienced. For each stimulation, only a single visual object image was shown, and the stimulation was triggered manually several seconds after the patient started to look at the stimulus.

### 2.4 Face-selectivity: fMRI

The comprehensive methods (stimuli, stimulation procedures) used for this fMRI study were the same as those used in several previous studies (combined in a large-scale analysis in Rossion et al., 2012). The fMRI experiment took place in September 2014 that is 5 months later than the SEEG exploration.

#### 2.4.1 Stimuli

Four categories of stimuli were used: photographs of faces (F), cars (C), and their phase-scrambled versions: scrambled faces (SF) and scrambled cars (SC). The face condition consisted of 43 pictures of faces (22 females) cropped so that no external features (hair, etc.) were revealed. All faces were shown in frontal view (for all stimulus information, see Rossion and Caharel, 2011). They were inserted in a gray rectangular background. Similarly, the car condition consisted of 43 pictures of different cars in a full-front view also embedded in a gray background. The scrambled stimuli were made using a Fourier phase randomization procedure (FFT with phase replaced by the phase of a uniform noise) that yields images preserving the low-level properties of the original image (i.e., luminance, contrast, spectral energy, etc.), while completely degrading any category-related information. Pictures of faces/cars and the phase scrambled face/car pictures subtended equal shape, size and contrast against the background.

#### 2.4.2 Paradigm

MB performed 4 runs of 11 min duration each. In each run, there were 6 blocks of 18 s duration for each of the 4 types of stimuli. Blocks were separated by a baseline condition (cross fixation) of 9 s. In each block, 24 stimuli of the same condition were presented (750 ms per stimuli, no ISI) on a black background screen, with 2 or 3 consecutive repetitions of the exact same stimulus in each block (target trials in the one-back task). This gave a total amount of 144 stimuli per category per run. The stimuli and the fixation cross were presented centrally, but stimulus location varied randomly in x (6%) and in y (8%) directions at each presentation. This change in stimulus location was made so that specific elements of the non-scrambled face and car stimuli (e.g., the eyes or headlights) do not appear at the same location at each trial, as it would be the case for scrambled stimuli even without jittering position. The patient performed a one-back identity task (2 or 3 targets per block).

#### 2.4.3 Imaging acquisition parameters

Functional MR images of brain activity were collected using a 3T head scanner (Signa HDXT, GE Medical Systems, Milwaukee, WI) at the University Hospital of Nancy with repeated single-shot echo-planar imaging: echo time (TE) = 33 ms, flip angle (FA) = 77°, matrix size = 64 × 64, field of view (FOV) = 192 mm, slice thickness = 3 mm, repetition time (TR) = 2250 ms, 36 slices. A high-resolution anatomical volume of the whole brain was acquired using a T1-weighted sequence (resolution: 1 × 1 × 1 mm).

#### 2.4.4 Data analysis

The fMRI signal in the different conditions was compared using Brain Voyager QX (Version 2.8.0, Brain Innovation, Maastricht, The Netherlands). Preprocessing consisted of a linear trend removal for excluding scanner-related signal, a temporal high-pass filtering applied to remove temporal frequencies lower than three cycles per run, and a correction for small interscan head movements by a rigid body algorithm rotating and translating each functional volume in 3D space. Functional data (unsmoothed) were spatially aligned with the high-resolution anatomical volume which was previously aligned to the AC-PC plane (automatic co-registration in Brain Voyager QX, adjusted manually). Subsequently, the functional data were analyzed using a multiple regression model (General Linear Model; GLM) consisting of predictors, which corresponded to the particular experimental conditions of each experiment. The predictor time courses used were computed on the basis of a linear model of the relation between neural activity and hemodynamic response, assuming a rectangular neural response during phases of visual stimulation.

The contrast of interest was the conjunction contrast [(F-C) and (F-SF)]. This contrast was aimed at isolating the regions responding more to faces than non-faces objects, and for which this difference could not be accounted for by low-level visual cues (Rossion et al., 2012). A conservative statistical threshold (Bonferroni-corrected, p&0.05) was used to define face-sensitive areas corresponding to t-values above 4.93.

#### 2.4.5 Intracerebral contact localization.

The high-resolution T1 (aligned to the AC-PC plane) was fused with the postoperative CT-scan. The electrode contact coordinates were automatically extracted (MRI coordinates in the individual anatomy centered on the AC-PC plane). These electrode contact coordinates were then rendered in Brain Voyager software. The anatomical locations of relevant fMRI activations and intracerebral contacts were therefore assessed in the individual anatomy. Anatomical and functional volumes were also spatially normalized (Talairach and Tournoux, 1988), but only to determine Talairach coordinates of fMRI activations and intracerebral contacts.

### 2.5 Face-selectivity: intracerebral responses

We used a “frequency-tagging” or fast periodic visual stimulation (FPVS) approach with natural images to identify and to quantify intracerebral face-selective responses. This paradigm has been validated in several studies (Rossion et al., 2015; de Heering and Rossion, 2015; Retter and Rossion, 2016) and consists of presenting widely variable face stimuli at regular intervals, here, as every 5 stimuli, in a fast periodic train of variable nonface object images (usually 6 Hz) (Figure 2). Frequency-domain representation of the EEG recorded during stimulation separates common responses to faces and objects at 6 Hz and its harmonics from face-selective responses occurring at 6 Hz/5, 1.2 Hz and its harmonics. The technique is highly sensitive and largely free of low-level visual confounds (Rossion et al., 2015), making it ideal for intracerebral recordings. The comprehensive methods used here (stimuli, stimulation procedures) were the same as in a recently reported intracerebral group study (Jonas et al., 2016).

**Figure 2.**
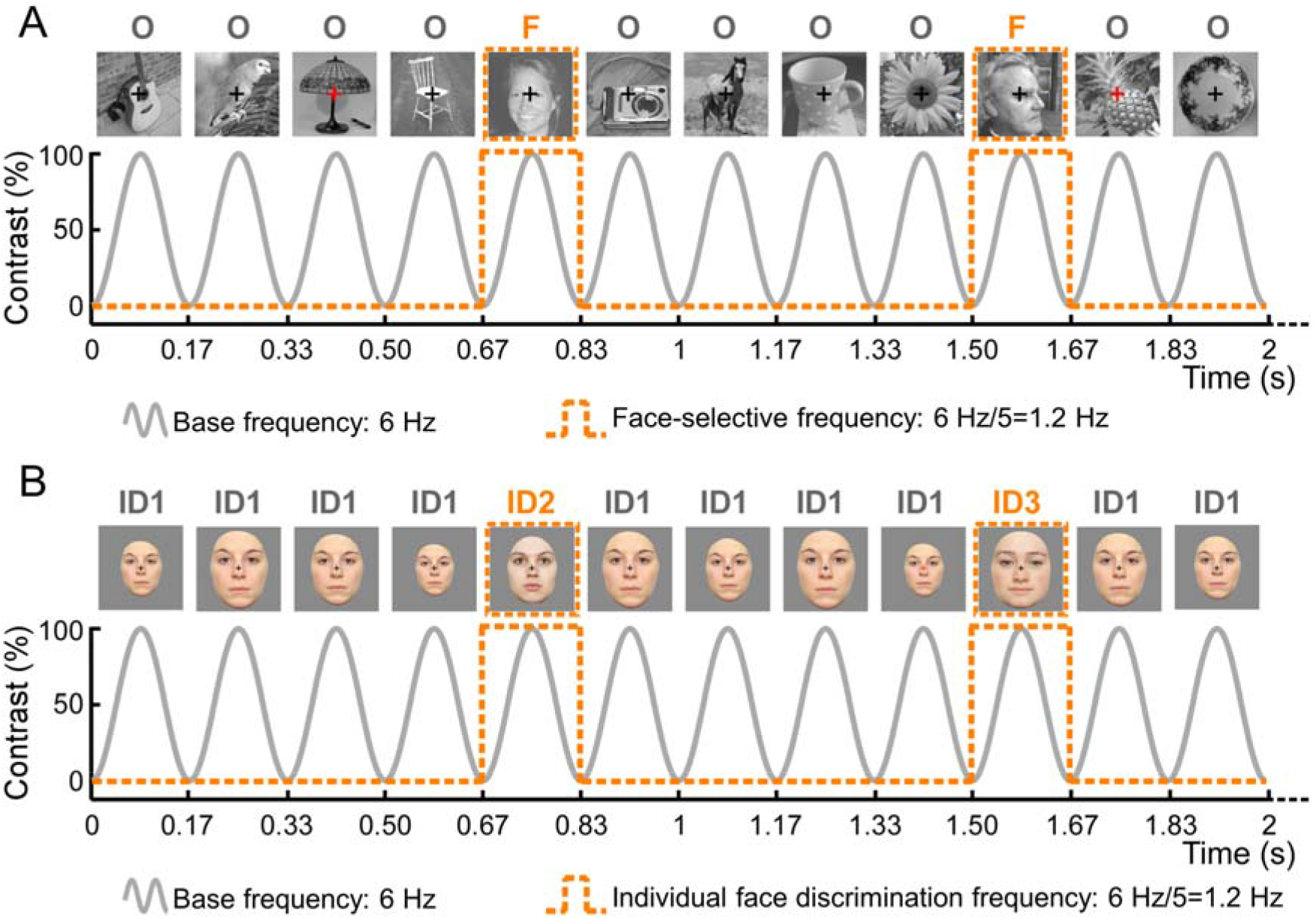
FPVS paradigms. **A**. Paradigm used to define face-selective neural activity (from Rossion et al., 2015). Natural object images are presented by sinusoidal contrast modulation at a rate of 6 stimuli per second (6 Hz). A different face image appears every 5 stimuli (i.e., 6 Hz/5=1.2 Hz). In these conditions, a significant electrophysiological response at 1.2 Hz and harmonics reflects a differential, i.e. selective, periodic response to faces (Rossion et al., 2015; de Heering and Rossion, 2015; Jonas et al., 2016). **B**. Paradigm measuring sensitivity to individual faces (from Liu-Shuang et al., 2014). Images of a randomly selected face identity are presented by sinusoidal contrast modulation at a rate of 6 Hz (ID1). Different face identities (ID2, ID3, etc.) appear every 5^th^ stimulus. Hence, face identity changes occurred at a rate of 6Hz/5=1.2Hz. Responses at this frequency and harmonics reflect individual face discrimination responses (Liu-Shuang et al., 2014; Liu-Shuang et al 2016).

#### 2.5.1 Stimuli

Two hundred grayscale natural images of various non-face objects (from 14 non-face categories: cats, dogs, horses, birds, flowers, fruits, vegetables, houseplants, phones, chairs, cameras, dishes, guitars, lamps) and 50 grayscale natural images of faces were used (see Figure 2A for examples of stimuli). Each image contained an unsegmented object or face near the center which differed in terms of size, viewpoint, lighting conditions and background. Images were equalized for mean pixel luminance and contrast.

#### 2.5.2 Experimental procedure

MB viewed 2 continuous sequences with highly variable natural images of objects presented at a fast rate of 6 Hz, with faces presented periodically as every 5th image (i.e., at 1.2 Hz = 6 Hz/5, see Figure 2A, see also Video 1 for an example of visual stimulation). All images were randomly selected from their respective categories. A sequence lasted 70 s: 66 s of stimulation at full-contrast flanked by 2 s of fade-in and fade-out, where contrast gradually increased or decreased, respectively (total duration of the experiment: 2×70 s=2 min 20 s). During the sequences, MB was instructed to fixate a small black cross which was presented continuously at the center of the stimuli and to detect brief (500 ms) color-changes (black to red) of this fixation-cross.

#### 2.5.3 Frequency domain processing

Segments of SEEG corresponding to stimulation sequences were extracted (74-second segments, -2 s to +72 s). The 74 s data segments were cropped to contain an integer number of 1.2 Hz cycles beginning 2 s after the onset of the sequence (right at the end of the fade-in period) until approximately 65 s, before stimulus fade-out (75 face cycles ≈ 63 s). The 2 sequences were averaged in the time domain. Subsequently, a Fast Fourier Transform (FFT) was applied to these averaged segments and amplitude spectra were extracted for all contacts.

#### 2.5.4 Face-selective responses

The FPVS approach used here allows identification and separation of two distinct types of responses (Rossion et al., 2015; Jonas et al., 2016): (1) a *general visual response* occurring at the base stimulation frequency (6 Hz) and its harmonics, as well as (2) a *face-selective response* at 1.2 Hz and its harmonics. Face-selective responses significantly above noise level at the face stimulation frequency (1.2 Hz) and its harmonics (2.4, 3.6 Hz, etc.) were determined by transforming the frequency spectra to Z-scores. The Z-scores were computed as the difference between amplitude at each frequency bin and the mean amplitude of the corresponding 48 surrounding bins (up to 25 bins on each side, i.e., 50 bins, but excluding the 2 bins directly adjacent to the bin of interest, i.e., 48 bins) divided by the standard deviation of amplitudes in the corresponding 48 surrounding bins. A contact was considered to be face-selective if a Z-score exceeded 3.1 (i.e., p<0.001 one-tailed: signal>noise) for at least one of the first 4 face-selective frequency harmonics (1.2, 2.4, 3.6 or 4.8 Hz; Jonas et al., 2016).

#### 2.5.5 Quantification of responses amplitude

Baseline-corrected amplitudes were computed as the difference between the amplitude at each frequency bin and the average of 48 corresponding surrounding bins (up to 25 bins on each side, i.e., 50 bins, but excluding the 2 bins directly adjacent to the bin of interest, i.e., 48 bins). The face-selective responses were then quantified at each face-selective contact as the sum of the baseline-subtracted amplitude across harmonics (Retter and Rossion, 2016). The range over which face and base frequency harmonics were summed was constrained by the highest significant harmonic (Z-score>3.1, p<0.001). In MB study, as in a previous intracerebral study of 28 subjects (Jonas et al., 2016), no significant face-selective responses were found above the 14th harmonic (i.e., 16.8 Hz). Face-selective responses were therefore quantified as the sum of the baseline-subtracted amplitudes at the face-selective frequency harmonics from the 1st until the 14th (1.2 Hz until 16.8 Hz), excluding the 5th and 10th harmonics (6 Hz and 12 Hz) that coincided with the base frequency. Signal-to-noise ratio (SNR) spectra were also calculated as the ratio between the amplitude at each frequency bin and the average of the corresponding 48 surrounding bins for display purposes and comparison across studies.

### 2.6 Visual discrimination of individual faces: intracerebral responses

We also used the FPVS approach to identify and to quantify intracerebral responses reflecting the visual discrimination of individual faces. The stimuli and experimental procedures are almost identical to the use of this paradigm on the scalp (Liu-Shang et al., 2014; Liu-Shang et al., 2016) but the methods are also detailed here for the first report of its application inside the brain.

#### 2.6.1 Stimuli

Full-front colored photographs of 25 male and 25 female faces with a neutral expression, taken under standardized conditions with respect to lighting, background, and distance from the camera were used (Figure 2B for examples of faces). External features such as hair and ears were cropped out and the isolated faces were put against a neutral grey background. Final images were resized to a height of 250 pixels (width: 186 ±11 pixels).

#### 2.6.2 Experimental procedure

MB viewed 2 continuous sequences of upright faces presented at a rate of 6 Hz (1 sequence with male faces and 1 with female faces). A sequence lasted 65 s: 60 s of stimulation at full-contrast ending with 5 s of fade-out, where contrast gradually decreased (total duration of the experiment: 4×65 s=4 min 20 s). In every sequence, the base face was a randomly selected face identity (within the 25 faces of one gender set) repeating throughout the sequence (e.g., identity 1, ID1). At fixed intervals of every 5th base face, a different facial identity (selected from the 25 remaining faces) was presented (ID2, ID3, ID4, etc.). Thus, a trial sequence contained face changes at a frequency of 6 Hz/5, i.e, 1.2 Hz (ID1-ID1-ID1-ID1-ID2-ID1-ID1-ID1-ID1-ID3-ID1-ID1-ID1-ID1-ID4, etc., see Figure 2B, see also Video 2). As a result, EEG amplitude at this frequency (1.2 Hz) and its harmonics (2.4 Hz, 3.6 Hz, etc.) was used as an index of the visual system’s discrimination of individual faces. To avoid confounding changes in face identity with changes with local pixel intensity changes (e.g., blue eyes vs. brown eyes), face size varied randomly between 74% and 120% in 2% steps at every 6 Hz stimulation cycle so that the low-level features of the faces did not overlap. During the sequences, MB fixated a small black cross presented continuously at the center of the stimuli and had to detect brief (500ms) color-changes of this fixation-cross (black to red).

#### 2.6.3 Frequency domain processing

Segments of SEEG corresponding to stimulation sequences were extracted (69-second segments, -2 s to +67 s). The 69 s data segments were cropped to contain an integer number of 1.2 Hz cycles beginning 2 seconds after the onset of the sequence until approximately 60 seconds, before stimulus fade-out (69 identity change cycles ≈ 58 s). The 2 sequences were averaged in the time-domain. Subsequently, FFT was applied to these averaged segments and amplitude spectra were extracted for all contacts.

#### 2.6.4 Individual face discrimination responses

The FPVS approach used here allows identifying and separating two distinct types of responses: (1) a *base response* occurring at the base stimulation frequency (6 Hz) and its harmonics, as well as (2) an *individual face discrimination responses* at 1.2 Hz and its harmonics. Individual face discrimination responses significantly above noise level at the face identity change frequency (1.2 Hz) and its harmonics (2.4, 3.6 Hz, etc.) were determined by transforming the frequency spectrums to Z-scores. Similarly to the face-localizer FPVS paradigm, the Z-scores were computed as the difference between amplitude at each frequency bin and the mean amplitude of the corresponding 48 surrounding bins divided by the standard deviation of amplitudes in the corresponding 48 surrounding bins. A contact was considered as showing individual face discrimination responses if a Z-score exceeded 3.1 (i.e., p<0.001 one-tailed: signal>noise) for at least one of the first 4 identity change frequency harmonics (1.2, 2.4, 3.6 or 4.8 Hz).

#### 2.6.5 Quantification of response amplitude

Baseline-corrected amplitudes were computed as the difference between the amplitude at each frequency bin and the average of 48 corresponding surrounding bins. The individual face discrimination responses were then quantified at each individual face discrimination contact as the sum of the baseline-corrected amplitude across harmonics (Retter and Rossion, 2016). The range over which face were summed was constrained by the highest significant harmonic across subjects (Z-score>3.1, p<0.001). In subject MB’s brain, no significant individual face discrimination responses were found above the 4th harmonic (i.e., 4.8 Hz). Individual face discrimination responses were therefore quantified as the sum of the baseline-corrected amplitudes at the identity change frequency harmonics from the 1st until the 4th (1.2 Hz until 4.8 Hz) (Liu-Shuang et al., 2016).

## 3. Results

### 3.1 Two intracerebral contacts within the face-selective right latFG

The fMRI face-localizer experiment identified typical face-selective activations of the “core” face network (right and left OFA, FFA, and posterior STS) and of the “extended” face network (right anterior temporal lobe) (Figures 3, Table 3). Contacts F1-F2 implanted in the latFG and in the adjacent occipito-temporal sulcus (Talairach coordinates: x: 34 to 37, y: -40, z: -17) were located within the right FFA (Figures 3 and 4A). FMRI activity in a 2-mm-radius ROI centered on the location of contacts was face-selective specifically for contacts F1 and F2 but not for adjacent contacts F3 and F4 (Figure 4B).

**Figure 3.**
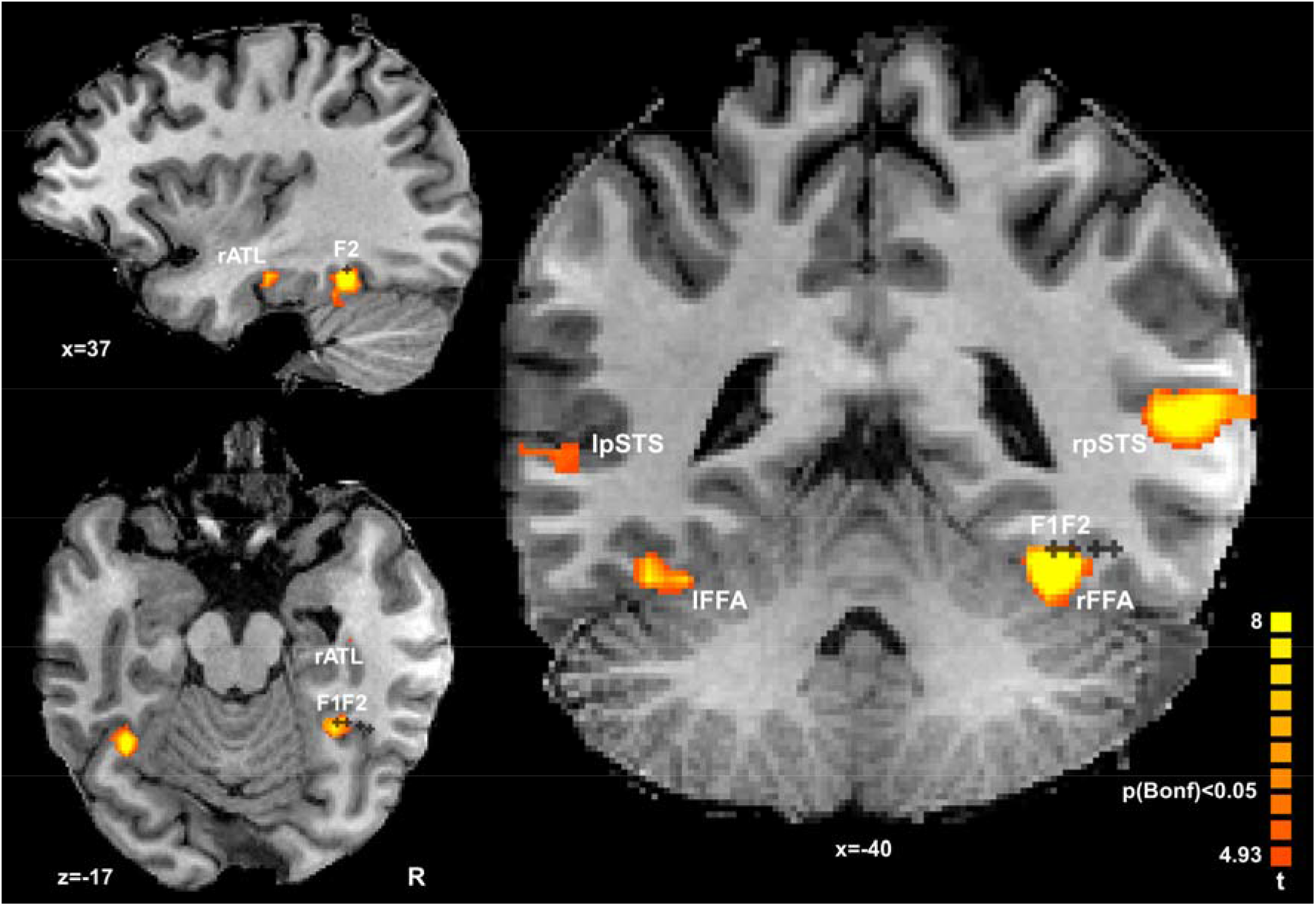
Two contacts F1 and F2 are located within the face-selective right latFG (“FFA”) defined in fMRI. FMRI revealed typical face-selective activations (conjunction contrast F-C and F-SF, p&0.05 Bonferroni-corrected): bilateral FFA, bilateral pSTS-faces, right ATL-faces and bilateral OFA (not shown here but see Table 3). Two contacts of electrode F (F1 and F2) were located within the right latFG-faces. Acronyms: ATL: anterior temporal lobe; FFA: fusiform face area; pSTS: posterior superior temporal sulcus.

**Table 3.**
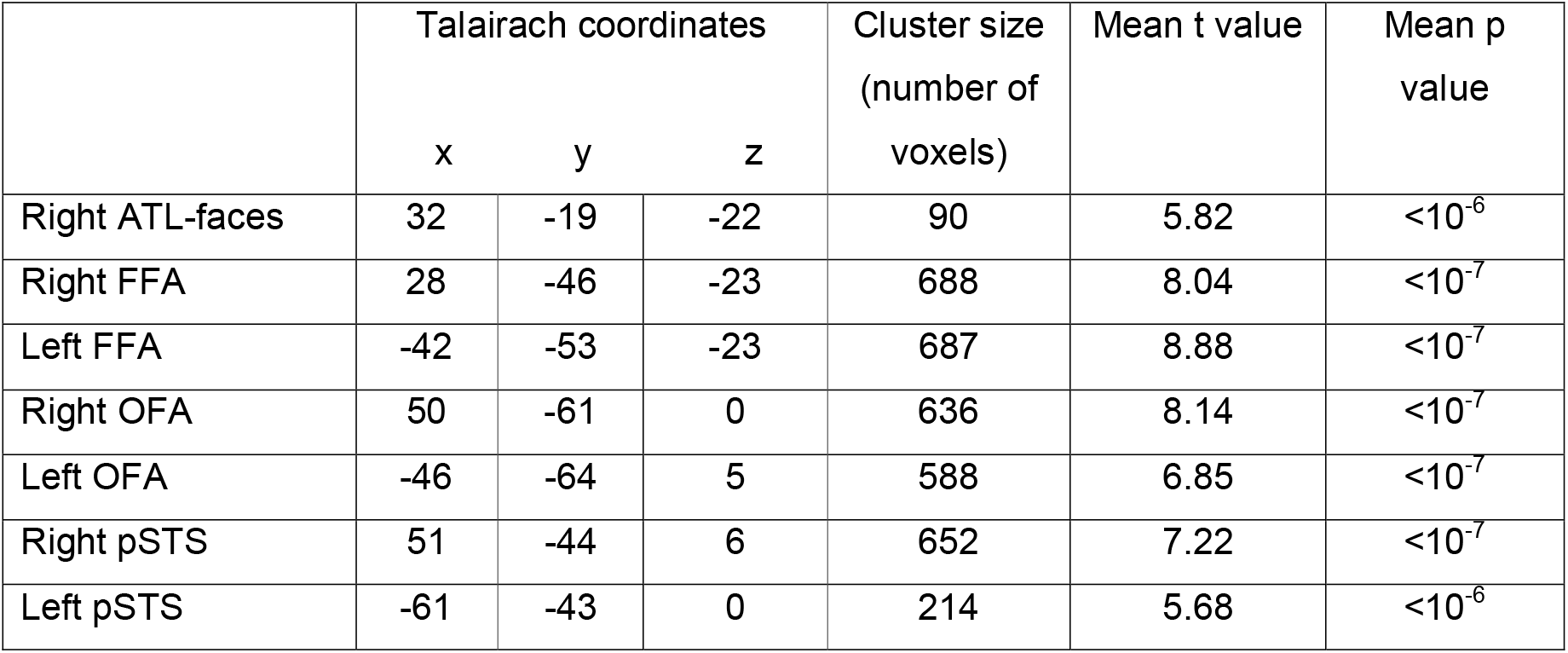
Talairach coordinates (center of mass), size, mean t and p values of face-selective areas identified in fMRI (conjunction contrast F–C and F–SF, p&0.05 Bonferroni-corrected). Acronyms: ATL: anterior temporal lobe; FFA: fusiform face area; OFA: occipital face area; pSTS: posterior superior temporal sulcus.

We used the FPVS approach to objectively (i.e., at experimentally-defined frequencies) identify intracerebral electrophysiological face-selective responses (Jonas et al., 2016). MB viewed sequences with highly variable natural images of objects presented at a fast rate of 6 Hz, with faces inserted every 5th image so that the face frequency was set at 1.2 Hz (6 Hz/5). In the frequency domain, responses at 1.2 Hz and harmonics (2.4 Hz, 3.6 Hz, etc.) reflect face-selective responses (Rossion et al., 2015; Retter and Rossion, 2016). On contacts F1 and F2 in the right FFA, we recorded large face-selective responses occurring exactly at 1.2 Hz and harmonics (Figure 4C). Significant face-selective responses were also found on 14 other contacts: contacts F3 and F4 located in the right occipito-temporal sulcus (Figure 4A), 2 contacts in the right middle temporal gyrus (lateral contacts of electrodes F), 4 contacts in the right inferior occipital gyrus (lateral contacts of electrode D), 2 contacts in the right anterior part of the collateral sulcus (electrode C) and 4 contacts in the ventral and medial part of the right occipital cortex (electrodes D and Q).

**Figure 4.**
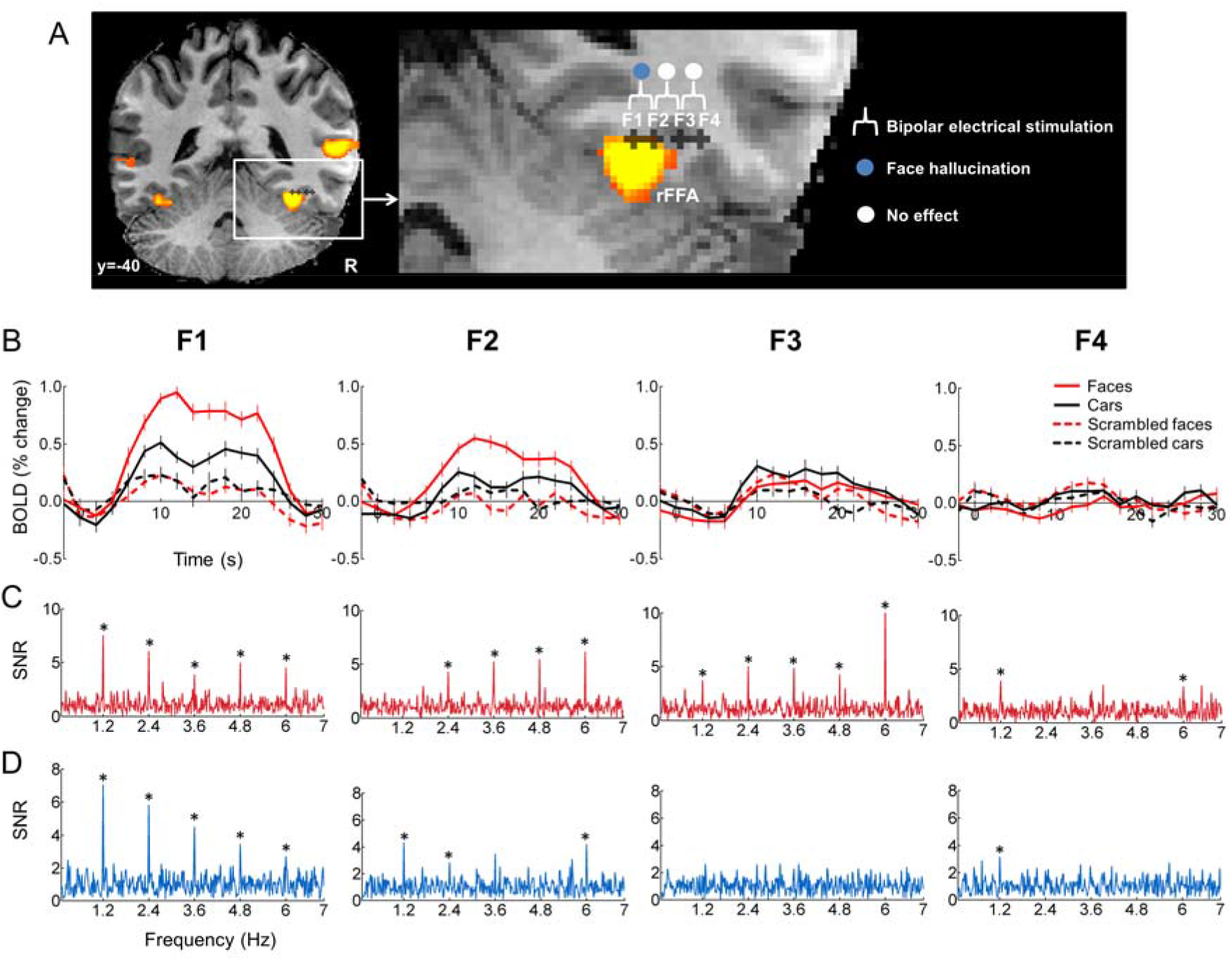
FMRI and electrophysiological measurements on contacts within and outside the right FFA. **A.** Locations of contacts F1, F2, F3 and F4. F1 and F2 were located within the right FFA while F3 and F4 were located just next to the FFA in the right OTS. Electrical stimulation of contacts F1 and F2 elicited individual face hallucinations. **B.** BOLD time courses recorded during the face-localizer fMRI experiment and extracted from a 2-mm-radius ROI centered on the location of contacts F1, F2, F3 and F4. **C.** Intracerebral responses recorded during the FPVS paradigm measuring face-selective activity. Significant face-selective responses occurring exactly at the face stimulation frequency (1.2 Hz and harmonics: 2.4, 3.6 and 4.8 Hz) were recorded on these 4 contacts with a maximum amplitude on contact F1 (see Figure 5A). *Indicates statistically significant responses (Z>3.1, p<0.001). **D.** Intracerebral responses recorded during the individual face discrimination paradigm. Significant individual face-discrimination responses exactly at the identity change frequency rate (1.2 Hz and harmonics) were recorded with a maximum amplitude at contact F1 (see Figure 5B). *Indicates statistically significant responses (Z>3.1, p<0.001).

We quantified the overall face-selective response amplitude on each contact (138 contacts in total) by summing the baseline-subtracted amplitudes over face-selective harmonics (sum of the 12 first face-selective frequency harmonics). Strikingly, contact F1 in the right FFA recorded the largest face-selective response amplitude among the 138 recording contacts implanted in MB’s brain (Figure 5A), followed by a series of contacts located adjacent to the FFA (F3), within the FFA (F2) and in the right inferior occipital gyrus (D9, D10, D11 and D12).

**Figure 5.**
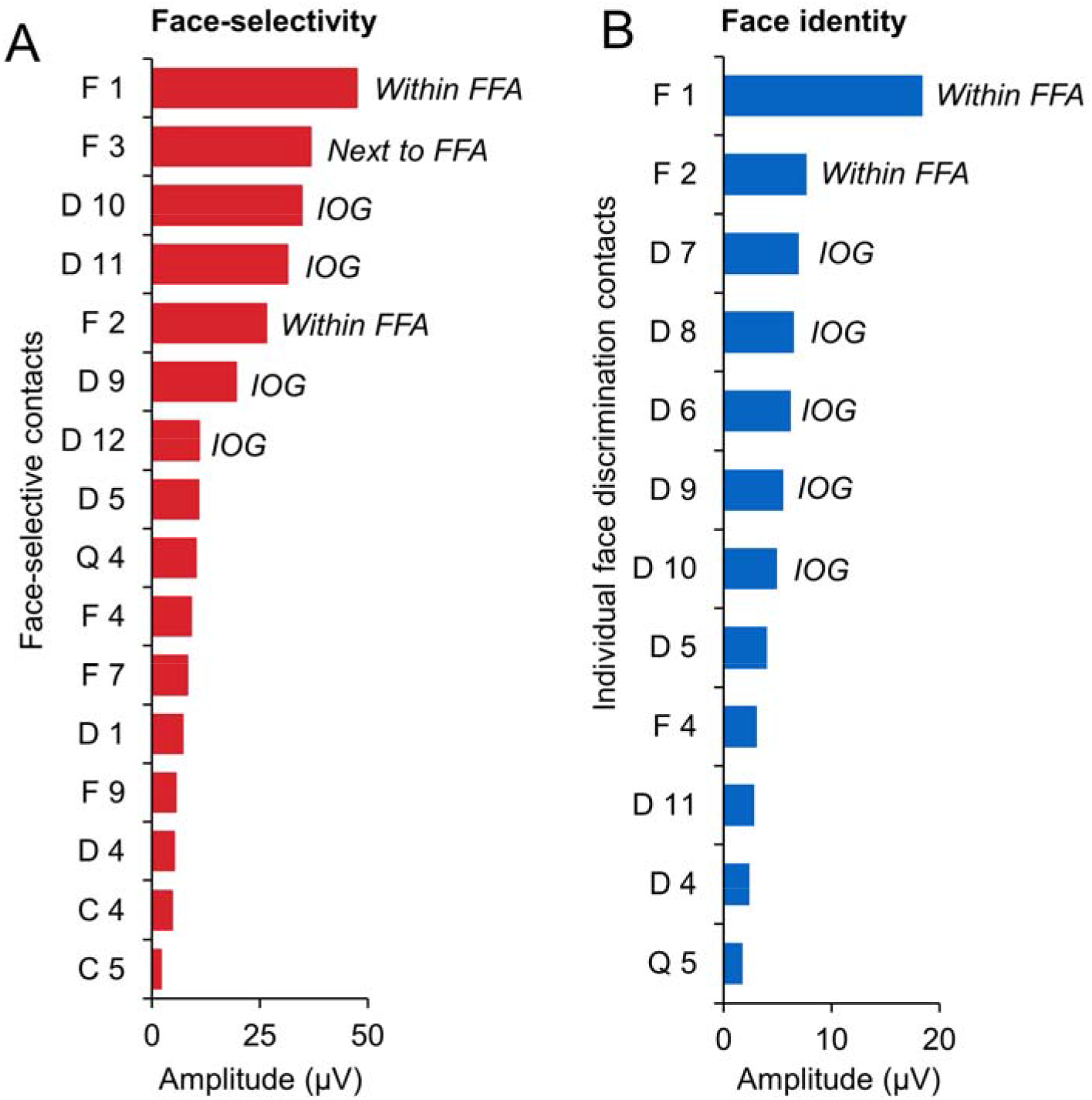
Quantification of the intracerebral response amplitude obtained with FPVS. **A.** Face-localizer FPVS paradigm. Among the 16 significant face-selective contacts, F1 located in the right FFA recorded the largest face-selective response amplitude. **B.** Face-identity FPVS paradigm. Among the 12 significant individual face-discrimination contacts, F1 located in the right FFA recorded the largest response amplitude. Acronyms: FFA: fusiform face area; IOG: inferior occipital gyrus.

Taken together, fMRI and intracerebral responses recorded during FPVS showed that 2 contacts (F1 and F2) were located within a face-selective region in the right latFG, and that of all contacts, F1 showed the largest face-selective response amplitude.

### 3.2 Stimulating the right latFG induces individual facial hallucinations (palinopsia) without distortions

Low intensity stimulations (1.2 mA) involving contacts F1-F2 located in the right latFG (Figure 4A) evoked hallucinations of individual face parts.

In the famous face recognition task in which 2 faces were successively presented during the time of the stimulation, MB reported seeing facial elements of the first famous face appropriately incorporated into the second famous face (stimulations 1 and 2). MB stated for stimulation 1: *“the photograph of Sarkozy* [former French president, first presented face during stimulation] was *transposed onto the other face* [second presented face during stimulation]”. Similarly, MB reported for stimulation 2: “I saw *these eyes* [indicating the first presented face during stimulation] *with this mouth and this mole* [indicating the second face during stimulation]” (see Video 3).

During stimulation of the same F1-F2 contacts, when MB was asked to passively look at a single face (a real face or a photograph of an unknown face), she reported hallucinations of individual facial elements incorporated appropriately into the face being perceived (stimulations 3, 4, 5 and 6). When MB was looking at the doctor’s face, she stated: “I saw *you with eyes and ears which were not yours”, “the ears were not yours, they were those of somebody else”* (stimulation 3, see Video 4). Stimulating these contacts when MB looked at a photograph of an unknown face, she stated: *“the eyes changed, the pupil took another form that is familiar to me”* (stimulation 6). MB reported that these facial parts were familiar to her but without being able to retrieve where and when she had already seen these features. She stated for example: *“they were not your eyes, they were the eyes of someone I had already seen, maybe coming from the images you showed me earlier”* (stimulation 3, see Video 4), *“It is something familiar but I can’t put a name on it; it is like when you meet someone you have already seen but without being able to retrieve his name”* (stimulation 6). Although MB was unable to remember whose face these features belonged to, these facial elements were always linked to individual familiar identities. Interestingly, the change in the percept concerned only part of the face, with some facial elements remaining intact. When asked if the nose changed, MB answered: “no” (stimulation 3); when asked if the eyes, the ears, the mouth, the hair changed, she answered: *“no, only the nose”* (stimulation 5). MB also spontaneously stated: *“only the eyes changed, the rest of the face did not change”* (stimulation 6).

Importantly, MB never reported any facial distortions and reproducibly stated that the facial structure was preserved *(“it* was *a normal face”* she said; when asked if the face was distorted, MB answered without hesitation: “no”). Once MB said *“your face* was *distorted”,* but it was a way to explain that the face changed because of the hallucinatory facial elements (stimulation 3, Video 4). Moreover, MB was always able to recognize the faces during the stimulation.

For the stimulations with non-face stimuli (active recognition of famous scenes and passive viewing of common objects), MB also reported individual face hallucinations (stimulations 7 and 8). During stimulation 7, MB was presented with famous scenes and asked to recognize them. MB reported seeing a pair of eyes looking at her, superimposed on the scene (Mont Saint-Michel in France, see Video 5). She stated: “I saw *the eyes that I saw earlier*”, “*they were not in place of something, they were superimposed on the scene*”. When asked where these eyes came from, she said: “I *don’t know which face but it* was *a face that I* saw *recently’.* During stimulation 8, MB was presented with a photograph of a car and was asked to passively looking at it. She reported seeing a whole face in the rear-view mirror. She stated: *“A face put itself in the rear-view mirror’, “It* was *something familiar”*. A third stimulation with active recognition of common object did not evoke any visual effect (stimulation 9). Other stimulations performed outside the face-selective right fusiform gyrus (50 stimulations in total) did not evoke similar effects or visual recognition impairments.

### 3.3 The stimulated region is sensitive to individual face identity

We tested MB with a FPVS experiment providing high SNR and behavior-free measures of the brain’s discriminative response to individual faces (Liu-Shuang et al., 2014; Liu-Shuang et al., 2016). MB viewed sequences with a randomly selected face identity presented at a fast rate of 6 Hz, with changes of stimulus size at every cycle and, critically, different face identities inserted every 5th image (identity change frequency =1.2 Hz, i.e. 6 Hz/5). In the frequency domain, responses at 1.2 Hz and harmonics (2.4 Hz, 3.6 Hz, etc.) reflect high-level individual face discrimination responses (i.e., abolished by inversion and contrast reversal of faces; and in acquired prosopagnosia; Liu-Shuang et al., 2014; Liu-Shuang et al., 2016).

At contact F1, we recorded large individual discrimination occurring exactly at 1.2 Hz and harmonics (Figure 4D). We also recorded clear, albeit smaller, responses on contact F2. Ten contacts also recorded individual discrimination responses, mainly in the right inferior occipital gyrus (D7, D8, D9, D10 and D11). We next quantified the individual face discrimination responses on each contact by summing the baseline-subtracted amplitudes over face-selective harmonics (sum of the 4 first harmonics). Contact F1 recorded, by far, the largest response amplitude (Figure 5B), followed by contact F2 and contacts in the right inferior occipital gyrus (D7, D8, D9, D10 and D11).

## 4. Discussion

We report a case of individual facial hallucination following the stimulation of the face-selective cortex of the right latFG. Two main characteristics make the reported hallucinations particularly original: (1) hallucinated face parts were linked to individual identities (i.e., to famous faces during the recognition task and to personally familiar faces during the passive face viewing task); (2) in the presence of a face stimulus, hallucinated faces parts were appropriately integrated within the perceived face. Thus, MB experienced confusion of facial identity perception by mixing different face parts coming from different identities in a coherent whole face. Moreover, these hallucinations were specific to the category of faces, as only face hallucinations were reported (regardless of the presence of face or non-face stimuli). Importantly, we also observed clear face-selective and individual face discrimination neural responses in the regions explored with intracerebral electrodes.

Puce et al. (1999) reported facial hallucinations by stimulating the face-selective cortex of the VOTC (isolated eyes, single or multiple faces). However, there was no report that these hallucinations belonged to particular individuals and patients were not looking at faces during stimulation. Moreover, these hallucinations followed stimulation of distributed sites along the postero-anterior axis of the VOTC. In contrast, MB hallucinations here followed the stimulation of a specific region not only highly selective to faces as defined in fMRI and SEEG, but also coding for differences between individual faces.

In two recent studies (Parvizi et al., 2012; Rangarajan et al., 2014), patients looking at faces during stimulation of the face-selective cortex in the right latFG reported “prosopometamorphopsia” (Bodamer, 1947; Hécaen and Angelergues, 1962; Mooney et al., 1965; Brust and Behrens, 1977; Ebata et al., 1991; Miwa and Kondo, 2007; Trojano et al., 2009; Hwang et al., 2012). Specifically, patients selfreported distortions of the perceived faces, involving the whole face or face parts (e.g., subject statements in Parvizi et al., 2012: *“Your face metamorphosed… your nose got saggy and went to the left”;* in Rangarajan et al. (2014)’s study, subject S3: *“Her nose looked different, larger”;* subject S4: *“Chin looks a little droopy”;* subject S5: *“It* was *almost like you were a cat”).* These reports are typical of prosopometamorphopsia, a condition in which faces may appear distorted, torn, warped, disfigured, or cartoonized, with, for example, face parts bulging, shrinking, dropping or floating. However, there is no hallucination or confusion between individual faces in such cases. The patient in Parvizi et al. (2012) stated that *“You almost look like somebody I’ve seen before, but somebody different”,* however, he also described severe distortions of the face *(“your nose got saggy and went to the left”, “it’s almost like the shape of your face, your features drooped”),* making unlikely that this transformed face corresponds to an existing individual. Moreover, subsequent observations described in Rangarajan et al. (2014) only mentioned prosopometamorphopsia. In contrast, MB never reported face distortions and stated that these hallucinations formed *“a normal face*”. In addition, brain-damaged patients experiencing prosopometamorphopsia are usually still able to recognize individual faces (Bodamer, 1947; Hécaen and Angelergues, 1962), which is line with patients’ reports during stimulation (Parvizi et al., 2012; Rangarajan et al., 2014).

A potential reason for the lack of individual face recognition impairment or hallucination following intracranial electrical stimulation of the right latFG in previous studies may be the size of the cortical region concerned. That is, electrical stimulation of the right latFG generating transient prosopometamorphopsia has been applied in previous studies through subdural electrodes posed on the cortical surface (“Electrocorticography”, ECoG), which are relatively large in size and distant from one another (i.e., electrodes of 2.3 mm in diameter with 5 to 10 mm inter-electrode spacing) and using high current intensities (2 to 8 mA). This approach contrasts with focal and low intensity electrical stimulation directly in the grey matter, as applied here with depth electrodes (SEEG, Bancaud and Talairach, 1973). Interestingly, in previous SEEG studies, focal intracerebral stimulation applied to other face-selective regions such as the right inferior occipital gyrus or the right anterior fusiform gyrus has sometimes elicited a transient impairment in individual face recognition (as observed in famous face recognition or face identity matching tasks), most often without face distortion (Jonas et al., 2012; Jonas et al., 2014; Jonas et al., 2015). However, we should make it clear from our experience that the observation of such phenomena following focal intracerebral stimulation within the latFG remains rare. A major limiting factor is obviously the positioning of the electrodes inside the brain, which follows strict clinical guidelines, and will often fall partly or completely outside of the critical cortical nodes for face perception. Hence, here the patient experienced recurrent face hallucinations following stimulation of a single electrode contact, while neighboring electrode contacts failed to elicit any effect (see also Jonas et al., 2012 for electrical stimulation at the edge of the right latFG/FFA without any effect on face perception).

MB’s visual experience corresponds to “palinopsia”, a condition in which there is persistence or recurrence of a visual image after the exciting object stimulus has been removed (Critchley 1951; Bender et al., 1968). Some types of palinopsia share a few characteristics with MB’s reports: recurrence of face images (“facial palinopsia”, all stimulations), recurrence appropriately integrated into the scene being perceived (all stimulations except number 7), recurrence of an object superimposed into comparable objects (“categorical incorporation”, e.g., eyes into a face, all stimulations with a face stimulus), recurrence after a short interval after exposure (“immediate form”, stimulations 1 and 2) and recurrence associated with a sense of familiarity, without being able to recall where or when the object has been seen before (stimulations 3 to 8) (Critchley, 1951; Kinsbourne and Warrington, 1963; Bender et al., 1968; Jacobs et al., 1972; Meadows and Munro, 1977; for a review see Gersztenkorn and Lee, 2015). For example, Meadows and Munro (1977) reported the famous case of a woman who, following a stroke of the right lingual and fusiform gyri, noticed that a Santa Claus beard became superimposed appropriately upon the faces of people at a Christmas party. Michel and Troost (1980) described two patients, who after a stroke of the right occipital lobe, reported that each person they saw had the face of someone they just had seen on television. Palinopsia including face hallucinations is usually reported following various lesions of the right occipito-temporo-parietal cortex and is usually associated with the recurrence of other object categories (Bender et al., 1968; Meadows and Munro, 1977; Michel and Troost, 1980; Maillot et al., 1993; Kupersmith et al., 1999). Our study shows that palinopsia restricted to faces can be evoked by stimulating a specific face-selective region of the right temporal lobe.

Here electrophysiological face-selective and face identity responses were recorded using a fast periodic visual stimulation approach providing objective, high SNR and quantifiable responses particularly useful for intracerebral recordings (e.g., Jonas et al., 2016). In the eloquent stimulation site, we recorded the largest face-selective and individual face discrimination responses among all the brain regions explored with intracerebral electrodes. Altogether, the generation of vivid hallucinations of individual faces with electrical stimulation and the recording of large individual face discrimination responses with FPVS point to the face-selective region in the right latFG as playing an active role in individual face representation. This statement is largely in agreement with findings of fMRI-adaptation studies, which have reported decreases to individual (unfamiliar) face repetition particularly in the face-selective cortex of the fusiform gyrus (e.g., Gauthier et al., 2000; Schiltz et al., 2006; Davies-Thompson et al., 2009; Ewbank et al., 2013; Xu and Biederman, 2010; Hermann et al., in press), with some studies showing a right hemispheric predominance of this effect (e.g., Winston et al., 2004; Mazard et al., 2006; Gilaie-Dotan and Malach, 2007). In addition, the disturbance reported by the patient here may be speculatively associated with a holistic/configural process (integration of multiple facial elements into a single representation of the whole face; e.g., Tanaka and Farah, 1993; Rossion, 2013). Indeed, hallucinated facial elements were appropriately incorporated within the face being perceived into a coherent face whole. Therefore, and although this is purely based on a subjective report, our observations may suggest a critical role of the right latFG in holistic/configural perception, a fundamental process for individual face recognition (Schiltz and Rossion, 2006; Rossion, 2013).

The co-localization between fMRI and intracranial electrophysiological responses is essential for electrical brain stimulation studies since it provides converging evidence of the functional specificity of the stimulated region. A spatial overlap between fMRI and electrophysiological face-selective responses has been shown in the latFG using transient stimuli (Puce et al., 1997; Parvizi et al., 2012; Jacques et al., 2016). Thanks to the FPVS approach here, we were not only able to show this co-localization, but also to quantify intracerebral responses and to rank brain regions according to their level of face-selectivity, this following a very short experiment time (e.g., 2 minutes here). One major problem in human electrical brain stimulation studies is the time constraint due to the clinical settings. In this context, FPVS appears to provide quick and reliable detection of the most important brain regions for a function, in order to primarily target them with electrical stimulation. This makes FPVS a tool of choice for subsequent electrical brain studies.

## 5. Conlusion

By stimulating the face-selective cortex in the right latFG, we evoked vivid percepts of individual faces. Along with intracerebral electrophysiological recordings during FPVS showing that this region is highly sensitive to individual faces, these results point out the active role of this region in individual face representation.

## Acknowledgments

We thank subject MB for their involvement in the study. We thank Joan Liu-Shuang for the FPVS individual face identity paradigm and Talia Retter for editing the manuscript. JJ and BR are supported by the Belgian National Fund for Scientific Research (FNRS). This work was supported by an European Research Council Grant (facessvep 284025).

